# Range-wide variation in local adaptation and phenotypic plasticity of fitness-related traits in *Fagus sylvatica* and their implications under climate change

**DOI:** 10.1101/513515

**Authors:** Homero Gárate-Escamilla, Arndt Hampe, Natalia Vizcaíno-Palomar, T. Matthew Robson, Marta Benito Garzón

## Abstract

**Aim:** To better understand and more realistically predict future species distribution ranges, it is critical to account for local adaptation and phenotypic plasticity in populations’ responses to climate. This is challenging because local adaptation and phenotypic plasticity are trait-dependent and traits co-vary along climatic gradients, with differential consequences for fitness. Our aim is to quantify local adaptation and phenotypic plasticity of vertical and radial growth, leaf flushing and survival across *Fagus sylvatica* range and to estimate each trait contribution to explain the species occurrence.

**Location:** Europe

**Time period:** 1995 – 2014; 2070

**Major taxa studied:** *Fagus sylvatica L.*

**Methods:** We used vertical and radial growth, flushing phenology and mortality of *Fagus sylvatica* L. recorded in BeechCOSTe52 (>150,000 trees). Firstly, we performed linear mixed-effect models that related trait variation and co-variation to local adaptation (related to the planted populations’ climatic origin) and phenotypic plasticity (accounting for the climate of the plantation), and we made spatial predictions under current and RCP 8.5 climates. Secondly, we combined spatial trait predictions in a linear model to explain the occurrence of the species.

**Results:** The contribution of plasticity to intra-specific trait variation is always higher than that of local adaptation, suggesting that the species is less sensitive to climate change than expected; different traits constrain beech’s distribution in different parts of its range: the northernmost edge is mainly delimited by flushing phenology (mostly driven by photoperiod and temperature), the southern edge by mortality (mainly driven by intolerance to drought), and the eastern edge is characterised by decreasing radial growth (mainly shaped by precipitation-related variables in our model); considering trait co-variation improved single-trait predictions.

**Main conclusions:** Population responses to climate across large geographical gradients are trait-dependent, indicating that multi-trait combinations are needed to understand species’ sensitivity to climate change and its variation across distribution ranges.

## 1. INTRODUCTION

Climate change is having a major impact on the structure, composition and distribution of forests worldwide (Trumbore, Brando, & Hartmann, 2015). Accordingly, numerous models have projected significant range shifts of forest tree species towards higher latitudes and elevations (Urban et al., 2016). However, to date, the two most important processes in the response of tree populations to a rapidly changing climate, local adaptation and phenotypic plasticity (Aitken, Yeaman, Holliday, Wang, & Curtis-Mclane, 2008; Savolainen, Pyhäjärvi, & Knürr, 2007), are not systematically considered by species distribution models (but see Duputié, Rutschmann, Ronce, & Chuine, 2015; Richardson, Chaney, Shaw, & Still, 2017; Valladares et al., 2014). Phenotypic plasticity enables a given genotype to express different phenotypes in response to changing environments, while local adaptation produces new genotypes with a greater ability to cope with the new environment. The two mechanisms are ubiquitous in natural populations, although their respective importance is considered to vary extensively through time and across species ranges (Des Roches et al., 2018; Reich et al., 2016). To better understand and more realistically predict future species distribution ranges, it is therefore critical to identify and quantify the respective importance of local adaptation and phenotypic plasticity in the response of local populations to a changing climate.

From an ecological perspective, fitness can be associated with several phenotypic traits which directly affect survival and reproduction, creating a fitness landscape (Laughlin, 2018) that allows them to be used to bound species ranges (Benito-Garzón, Ruiz-Benito, & Zavala, 2013; Stahl, Reu, & Wirth, 2014). Fitness-related traits vary across large geographical gradients, mainly depending on how natural selection drove differences among populations in the past. For instance, tree height is generally greatest at the core of a species range and decreases towards its margins (Pedlar & McKenney, 2017). Climate-driven mortality commonly increases towards the driest part of a species range, which is related to drought-induced stress conditions (Benito Garzón et al., 2018). The onset of flushing phenology tends to be delayed towards high latitudes (Duputié et al., 2015) as a consequence of genetic adaptation to late frost and fluctuating photoperiod (Way & Montgomery, 2015). Moreover, traits tend to co-vary across climatic gradients (Laughlin & Messier, 2015). A conspicuous example is the demographic compensation found between survival and growth near range margins (Benito-Garzón et al., 2013; Doak & Morris, 2010; Peterson, Doak, & Morris, 2018). New climatic conditions can result in maladaptation of some populations, which may change intra-specific patterns of trait variation and co-variation across geographical gradients, and eventually, species ranges. For example, increasing temperatures at high-latitude or high-elevation range margins are likely to produce higher growth rates, but they can also induce higher mortality owing to late frosts (Delpierre, Guillemot, Dufrêne, Cecchini, & Nicolas, 2017; Vitasse, Lenz, Hoch, & Korner, 2014). Hence, species ranges are likely to be delimited by the interaction of multiple traits and their responses across environmental gradients (Benito-Garzón et al., 2013; Enquist et al., 2015; Stahl et al., 2014).

Common gardens or provenance tests provide us with the necessary experiments to quantify phenotypic plasticity and local adaptation of fitness-related traits in response to climate (Mátyás, 1999). Models based on reaction norms of phenotypic traits using measurements recorded in common gardens show that: (i) geographic variation in populations’ responses to climate is more strongly based on phenotypic plasticity than on local (Benito Garzón, Robson & Hampe, *Under Review*); (ii) phenotypic variation can strongly differ among traits, in particular for tree survival, growth, and flushing phenology - traits that are directly related to fitness and typically measured in common gardens (Benito Garzón, Alía, Robson, & Zavala, 2011; Duputié et al., 2015; Richardson et al., 2017; Valladares et al., 2014); (iii) as a consequence, predictions of future species ranges are likely to be strongly influenced by the combined response of different fitness-related traits to climate (Laughlin, 2018), but this structured combination of intra-specific multi-trait variation defining species ranges has not been explored with empirical data.

*Fagus sylvatica* L. (European beech, henceforth “beech”) is a widely distributed deciduous broadleaved temperate tree. In some parts of its range, beech has a late flushing strategy to avoid late frosts, which has a generally detrimental effect on tree growth (Delpierre et al., 2017; Gömöry & Paule, 2011; Robson, Rasztovits, Aphalo, Alia, & Aranda, 2013). Beech is currently expanding at its northern distribution edge, whereas it experiences drought-induced radial growth decline and increasing mortality at its southern edge (Farahat & Linderholm, 2018; Stojnic et al., 2018). The extent to which this pattern will continue in the future depends on how the combination of several fitness-related traits will influence the species’ response to new climates.

Here, we propose a new modeling approach that quantifies local adaptation and phenotypic plasticity of four major phenotypic traits related to fitness (vertical and radial growth, survival, and flushing phenology) and their interactions, to delimit species ranges under current and future climates. To this purpose, we use the phenotypic measurements recorded in the BeechCOSTe52 database (Robson, Benito Garzón, & BeechCOSTe52 database consortium, 2018), the largest network of common gardens for forest trees in Europe, covering virtually the entire distribution range of the species. Our specific objectives are: (i) to quantify range-wide patterns of phenotypic plasticity and local adaptation in growth, survival and flushing phenology; (ii) to identify interactions among the different traits and the extent of their geographical variation in local adaptation and phenotypic plasticity; (iii) to discuss how these fitness-related traits delimit species ranges, and (iv) to better understand species ranges under new climate scenarios and the role of trait variation in shaping the future species range.

## 2. MATERIAL AND METHODS

We calibrated two types of linear mixed-effect models using a combination of trait measurements from common gardens where seeds coming from provenances from different origins have been planted (provenances) and of environmental variables that we obtained for these common gardens and provenances. The first model type (one-trait models) used single traits as response variables and environmental data as explanatory variables. The second model type (two-trait models) added a second trait as co-variate, which allowed the interaction of both traits to be accounted for in the model. Finally, to quantitatively estimate the contribution of each trait to explain beech range, we performed a binomial model using the occurrence of the species as response variable (presence/absence) and the spatial predictions of all traits as explanatory variables.

### 2.1. Trait measurements

We analyzed total tree height (vertical growth), diameter at breast height (DBH; radial growth), survival and flushing phenology measured on a total of 153,711 individual beech trees that originated from seeds collected from 205 populations (hereafter referred to as “provenances”) across Europe and planted at 38 common gardens (hereafter “trials”) (Table 1; Supporting Information Figure S1.1, Appendix S1). Briefly, the seeds were germinated in greenhouses and planted in the trials at an age of two years. Plantations were carried out in two consecutive campaigns, the first campaign (comprising 14 trials) in 1995 and the second one (comprising 24 trials) in 1998 (see more details in Robson et al., 2018). Survival was recorded as individual tree survival. Leaf flushing was transformed from observational stage score data (qualitative measurements that slightly differ among trials) to Julian days by adjusting flushing stages for each tree with the Weibull function by trial (Robson et al., 2011, 2013).

**Table 1.** The extent of data from the BeechCOSTe52 database used for modelling. Measurements: total number of measurements; Trees: total number of individual trees; Trials: total number of trials; Provenances: total number of provenances, Age: the age at which the trees were measured. Columns indicate sample sizes for the traits: tree height (Height), diameter at breast height (DBH), survival and leaf flushing that were used in the one-trait models, as well as for the combined height-DBH, and height-leaf flushing records that were used in the two-trait models.

|  | **Height** | **DBH** | **Survival** | **Leaf flushing** | **H-DBH** | **H-Leaf flushing** |
| --- | --- | --- | --- | --- | --- | --- |
| **Measurements** | 203 105 | 34 237 | 41 309 | 7 863 | 34 237 | 12 087 |
| **Trees** | 108 415 | 31 339 | 37 433 | 7 863 | 31 339 | 10 634 |
| **Trials** | 36 | 19 | 7 | 7 | 19 | 6 |
| **Provenances** | 205 | 186 | 114 | 62 | 186 | 150 |
| **Age** | 2 to 15 | 8 to 15 | 2 to 6 | 12 | 8 to 15 | 6, 9, 11, 12, 15 |

### 2.2. Environmental data

We used the EuMedClim database that gathers climatic information from 1901 to 2014 gridded at 1km (Fréjaville & Benito Garzón, 2018). The climate of the provenances was averaged for the period from 1901 to 1990, with the rationale that the seeds planted in the common gardens stemmed from trees growing during that period (Leites, Robinson, Rehfeldt, Marshall, & Crookston, 2012). To characterize the climate of the common gardens, we calculated average values for the period between the date of planting (either 1995 or 1998) and the year of measurement of each trait for 21 climate variables (Supporting Information Table S1.1, Appendix S1). In addition, we used the latitude and longitude of the provenance and of the trial as proxies for the photoperiod and continentality, respectively (used in our flushing phenology models).

Phenotypic predictions under future climates were performed using the representative concentration pathway (RCP 8.5) in GISS-E2-R from WorldClim (http://www.worldclim.org/cmip5_30s) for 2070. We deliberately chose only this pessimistic scenario because for long-lived organisms such as forest trees it makes little difference whether the projected situation will be reached in 2070 or some decades later.

### 2.3. Statistical analysis

#### 2.3.1. Environmental variable selection

To avoid co-linearity and reduce the number of environmental variables to use in models, we performed two principal component analyses (PCA), one for the climate variables related to the provenance site and one for the climate variables related to the trial site. For tree height, DBH and survival, we considered 21 variables for the provenance and 21 variables for the trial (Supporting Information Figure S1.2, Appendix S1); whereas for leaf flushing, we only included the temperature-related variables as well as latitude and longitude (a total of 20 variables), because leaf flushing is known to be mainly driven by them (Basler & Körner, 2014).

The retained variables after the PCA screening were combined in models containing one variable to characterize the climate of the provenance and one variable to characterize the climate of the trial (Supporting Information Table S1.2, Appendix S1). We selected the combination of environmental variables for each trait that minimize the AIC (Supporting Information Table S1.2, Appendix S1).

#### 2.3.2. One-trait and two-trait mixed-effect models

We used linear mixed-effect models to analyze the response of individual traits (one-trait models) and the co-variation between two traits (two-trait models) to climate. We included the climate at the provenance and the trial site as previously selected (Supplementary Table 1), the age of trees, and for the leaf flushing model also latitude and longitude as fixed effects. The trial, blocks nested within the trial and trees nested within block and trial, were included as random effects to control for differences among sites and for repeated measurements of the same trees. The random effect of the provenance was also included in the model. The common form of the one-trait model was: 

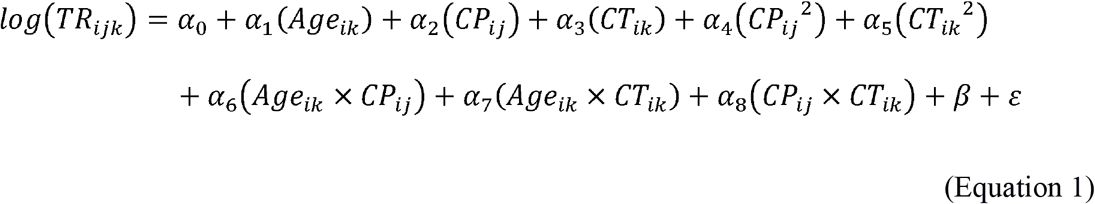

Where TR = trait response of the *i*^*th*^ individual of the *j*^*th*^ provenance in the *k*^*th*^ trial; Age = tree age of the *i*^*th*^ individual in the *k*^*th*^ trial; CP = climate at the provenance site of the *i*^*th*^ individual of the *j*^*th*^ provenance; CT = climate at the trial site of the *i*^*th*^ individual in the *k*^*th*^ trial; β = random effects and ε = residuals. In addition, the model included the following interaction terms: Age and CP, Age and CT, and CP and CT.

We analyzed trait co-variation across the species range by adding two specific traits of interest in the same model. The common form of the two-trait model was: 
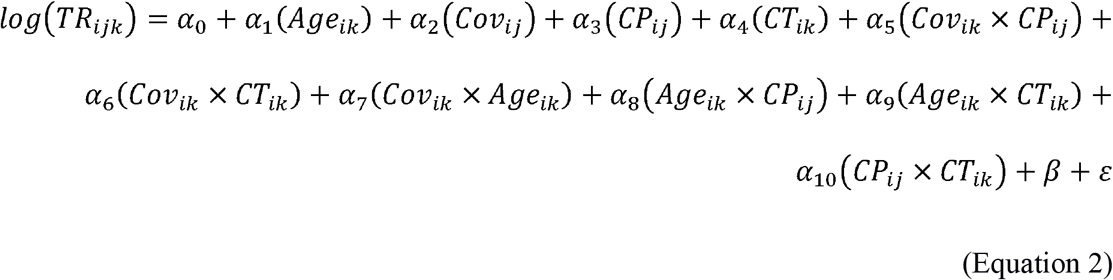

Where TR = trait response of the *i*^*th*^ individual of the *j*^*th*^ provenance in the *k*^*th*^ trial; Age = tree age of the *i*^*th*^ individual in the *k*^*th*^ trial; Cov = trait co-variate of the *i*^*th*^ individual in the *k*^*th*^ trial; CP = climate at the provenance site of the *i*^*th*^ individual of the *j*^*th*^ provenance; CT = climate at the trial site of the *i*^*th*^ individual in the *k*^*th*^ trial; β = random effects and ε = residuals. In addition, the model included the following interaction terms: Cov and CP, Cov and CT, Cov and Age, Age and CP, Age and CT, and CP and CT.

The one-trait and two-trait models for vertical and radial growth and leaf flushing were fitted with the ‘lmer’ function, while the one-trait model for survival was fitted with the ‘glmer’ function to accommodate logistic regressions (binomial family) in the analysis. We implemented a stepwise-model procedure with four main steps to choose the best supported model (Akaike, 1992): (i) we fitted saturated models that included all the variables in the fixed part of the model; (ii) we chose the optimal random component of the model by comparing the battery of models using restricted maximum likelihood (REML), and selected the best model using the Akaike information criterion (AIC) with criteria delta <2 (Mazerolle, 2006); (iii) we compared the battery of models using maximum likelihood (ML) and selected the optimal fixed component using the AIC criterion; (iv) we combined the best optimal random and fixed component previously selected and adjusted them using REML to obtain the best performing model. All model fits were done using the package ‘lme4’ (Bates et al., 2018).

For the best supported models, we visually analyzed the interactions of vertical growth, radial growth, survival and leaf flushing with the environment (one-trait models) and between traits (between the response and co-variate variable, i.e. the two-trait models). To do so, tree age was fixed to 12 years for the radial and vertical growth and leaf flushing models and to 6 years for the survival model. Mathematical interactions in one-trait models (CP x CT in equation 1) represent the differences in trait values that can be attributed to the provenance (interpretable as local adaptation) and those that can be attributed to the trial (interpretable as phenotypic plasticity). Mathematical interactions in two-trait models (Cov x CT in equation 2) represent the differences in trait values that can be attributed to a second trait that co-varies across the species range with the first trait, mediated by the climate of the trial (representing phenotypic plasticity). Unfortunately, survival could not be included in the two-trait models because there were insufficient measurements shared with other traits in the same trials.

We estimated the percentage of the variance explained by the model attributed to the fixed effects alone (marginal *R*^2^) and attributed to the fixed and random effects together (conditional *R*^2^). We measured the generalization capacity (Pearson correlation) of the model using cross-validation (64% of the data used for calibration and the remaining 34% for validation).

#### 2.3.3. Spatial predictions

We made spatial predictions for each trait across the species range for current and future climatic conditions using the ‘raster’ package (Hijmans et al., 2017). For the prediction of current and future trait variation, the climate variable for provenance was represented by the average climate over the period from 1901 to 1990. The climate of the trial was set as the average climate from 2000 to 2014, for current trait predictions, and to 2070 for future predictions. For two-trait models, the predicted values of the co-variate (DBH and leaf flushing) in the present were used to estimate the predictions of vertical growth in the future. We calculated the spatial difference between the future and the current conditions (future values minus current values) to illustrate the amount of change that traits can accommodate. All spatial predictions of traits were delimited within the distribution range of the species (EUFORGEN, 2009).

#### 2.3.4. Quantification of the trait contribution to delimit the range of beech

We regressed the occurrence (presence/absence) of the species (EUFORGEN, 2009) against the trait values obtained by the one-trait models using the ‘glm’ function to accommodate logistic regressions (binomial family). The equation takes the form: 
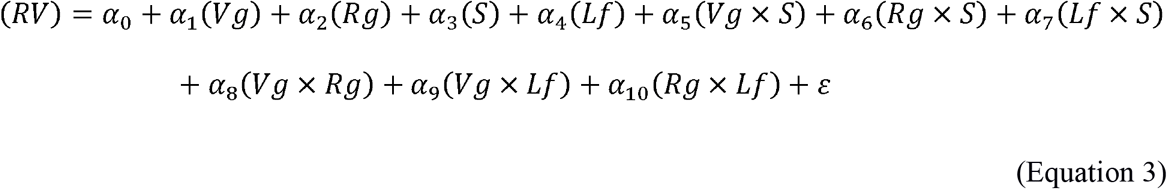

Where RV = presence/absence of beech; Vg = vertical growth; Rg = radial growth; S = survival; Lf = leaf flushing; ε = residuals. In addition, the model included all possible pairwise linear interactions of the included traits. The total deviance explained by the model was calculated using the function ‘Dsquared’ of the package ‘modEvA’ (Barbosa, Brown, & Real, 2014). Then, we performed an analysis of variance (ANOVA) of the model to obtain trait and trait interaction deviances to estimate the percentage of the variance attributable to each trait.

All the models where performed with the R statistical framework version 3.2.0 (R Development Core Team, 2015).

## 3. RESULTS

### 3.1. Environmental variables selection

The two PCA performed (provenance PCA and trial PCA) revealed two groups of variables, one related with temperature and another more related with precipitation (Supporting Information Figure S1.2, Appendix S1). The two most important axes of the provenance PCA explained 53.52 and 24.03% of the total variance, and those of the trial PCA explained 38.93 and 24.19% (Supporting Information Figure S1.2, Appendix S1). We to retain the following variables for tree growth and survival: BIO1, BIO5, BIO6, BIO12, BIO13, BIO14, Pet Mean and Pet Max. For the leaf flushing models, we retained BIO1, BIO5, BIO6, MTdjf, MTmam, MTjja, Mtson, and Mtdjfmam in addition to latitude and longitude.

### 3.2. One-trait and two-trait models

According to the best supported models (Table 2), the most important variable related to the climate at the provenance for vertical growth, radial growth and survival was maximal potential evapotranspiration (Pet Max). The most important variables related to climate at the trials were precipitation of the wettest month (BIO13) for vertical growth, annual precipitation (BIO12) for radial growth, and precipitation of the driest month (BIO14) for survival. In the case of leaf flushing, the mean temperature of December, January and February (MTdjf) was the most important climate variable for both the provenance and the trial site. The latitude of the provenance and the trial and the longitude of the trial were also significant in the leaf flushing model (see Supporting Information Table S1.3, Appendix S1 for detailed statistics on the models). We observed significant interactions between the climate of the trial and that of the provenance in all models (Table 2; Supporting Information Table S1.3, Appendix S1).

**Table 2.**
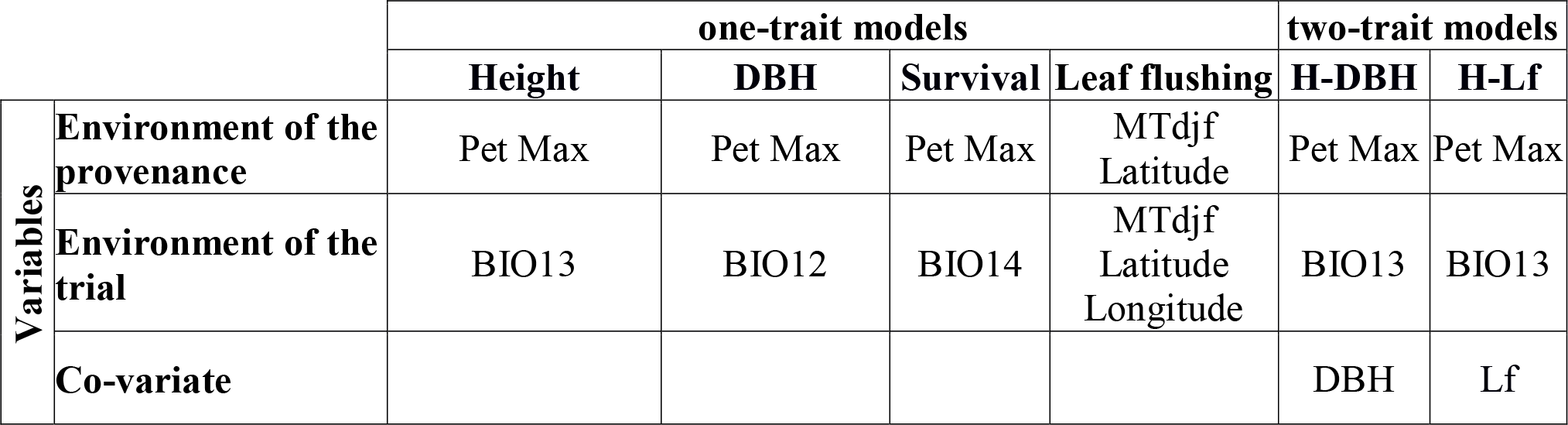
Summary of the variables included in the final best-supported models (one- and two-trait) for each trait analyzed. Environmental variables selected for the provenances and the trials for the one-trait models of height, DBH, survival and flushing, and for the two-trait models of height-DBH and height-leaf flushing. H: height; DBH: diameter at breast height; Lf: leaf flushing; Pet Max: maximal monthly potential evapotranspiration; BIO12: annual precipitation; BIO13: precipitation of wettest month; BIO14: precipitation of driest month; MTdjf: mean temperature of December, January and February; Co-variate: trait covariate.

The capacity for generalization from the models (Pearson correlation coefficients) was high: between 0.53 for radial growth and 0.73 for leaf flushing. The marginal *R*^2^ ranged from 18% for the survival model to 57% for the vertical growth model, while the conditional *R*^2^ ranged from 40% for the survival model to 98% for the radial growth model (Supporting Information Table S1.3, Appendix S1).

The significance of the fixed and random effects in the one-trait models was positively affected (i.e., estimates were higher) by the addition of a second trait (Supporting Information Table S1.4, Appendix S1). Furthermore, the co-variates and their interactions with the climate variables of the trials were also significant in the two-trait models (Supporting Information Table S1.4, Appendix S1). The capacity to generalize from the two-trait models was high: 0.76 for the vertical growth-radial growth model and 0.77 for the vertical growth-leaf flushing model (Supporting Information Table S1.4, Appendix S1). The marginal *R*^2^ was 62% in the vertical growth-radial growth model and 47% in the vertical growth-leaf flushing model, while the conditional *R*^2^ was 95% in the vertical growth-radial growth model and 99% in the vertical growth-leaf flushing model (Supporting Information Table S1.4, Appendix S1).

### 3.3. Spatial patterns of phenotypic trait variation from one-trait models

Spatial predictions showed differences in phenotypic trait variation among traits (Figure 1, maps) and the interaction graphs permitted the way that plasticity and local adaptation shape these differences to be visualized (Figure 1, interaction graphs).

**Fig 1.**
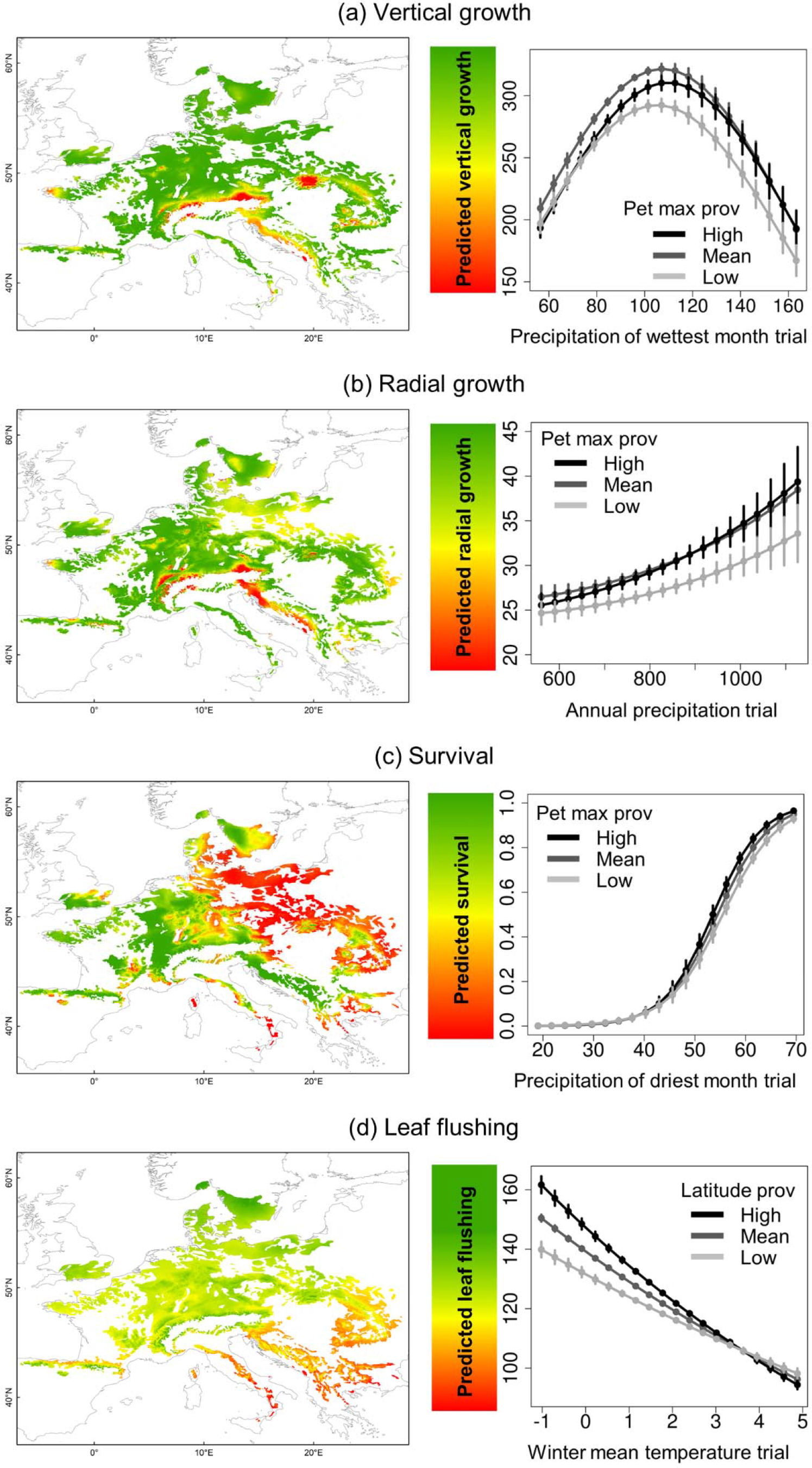
Spatial projections for (a) vertical growth (cm), (b) radial growth (mm), (c) survival (probability) and (d) leaf flushing (Julian days) generated using one-trait models (maps on the left), and corresponding graphs of interactions between the best environmental predictor variable across the trials divided according to environment at the provenance for each of the four traits (graphs on the right). Black, dark grey, and light grey lines represent high, medium and low values of the climatic variable of the provenances (as opposed to those of the trial, indicated on the x-axis). The vertical lines represent the confidence intervals. The maps display the trait projection for contemporary climate (inferred from 2000-2014 meteorological data) across the current species range. The color gradient depicts the clinal variation from low (red) to high (green) values of each trait. The values of the different traits are represented in the following way: vertical growth (cm), radial growth (mm), probability of survival (0 =dead, 1=alive) and leaf flushing (Julian days). Pet max prov: maximal monthly potential evapotranspiration at the provenance; Latitude prov: latitude of the provenance.

Vertical growth reached its maximum value at intermediate values of precipitation of the wettest month in the trials (Figure 1a, interaction graph). These largest trees were predicted to occur mostly over the northern and western part of the species range (Figure 1a, map). A signal of local adaptation was detected in our models and is shown by the interaction graph, where each line represents the response of provenances to high, intermediate and low levels of maximal potential evapotranspiration.

Predicted radial growth across the species range presented a similar pattern to that of vertical growth, with the lowest values in marginal populations, particularly at the southern margin (Figure 1b, map). High annual precipitation coincided with high growth rates (Figure 1b map), with a moderate signal of local adaptation in the form of some variation among provenances (Figure 1b, interaction graph).

The lowest survival rates were predicted towards the east and at some isolated points in the southernmost part of the range (Figure 1c, map). Survival increased towards those trials where precipitation is high in the driest month, with weak local adaptation indicated by very small–though statistically significant–differences among provenances (Figure 1c, interaction graph).

Earlier flushing was predicted towards the south-eastern part of the range (Fig 1d, map), with notable local adaptation indicated by large differences among provenances depending on the latitude of origin (Figure 1d, interaction graph). Differences in flushing date among provenances were particularly large in trials where the winter temperature is low (Figure 1d, interaction graph).

### 3.3. Patterns of phenotypic trait variation from two-trait models

Overall, models with a second trait as co-variate produced different results to those considering a single trait only. Predicted vertical growth was larger across the range when either radial growth (Figure 2a) or leaf flushing (Figure 2b) was included as a co-variate. Vertical growth increased with radial growth and precipitation (Figure 2a) and decreased in those regions where leaf flushing was predicted to be late in the year (which corresponded mainly to the northern part of the range) (Figure 2b).

**Fig 2.**
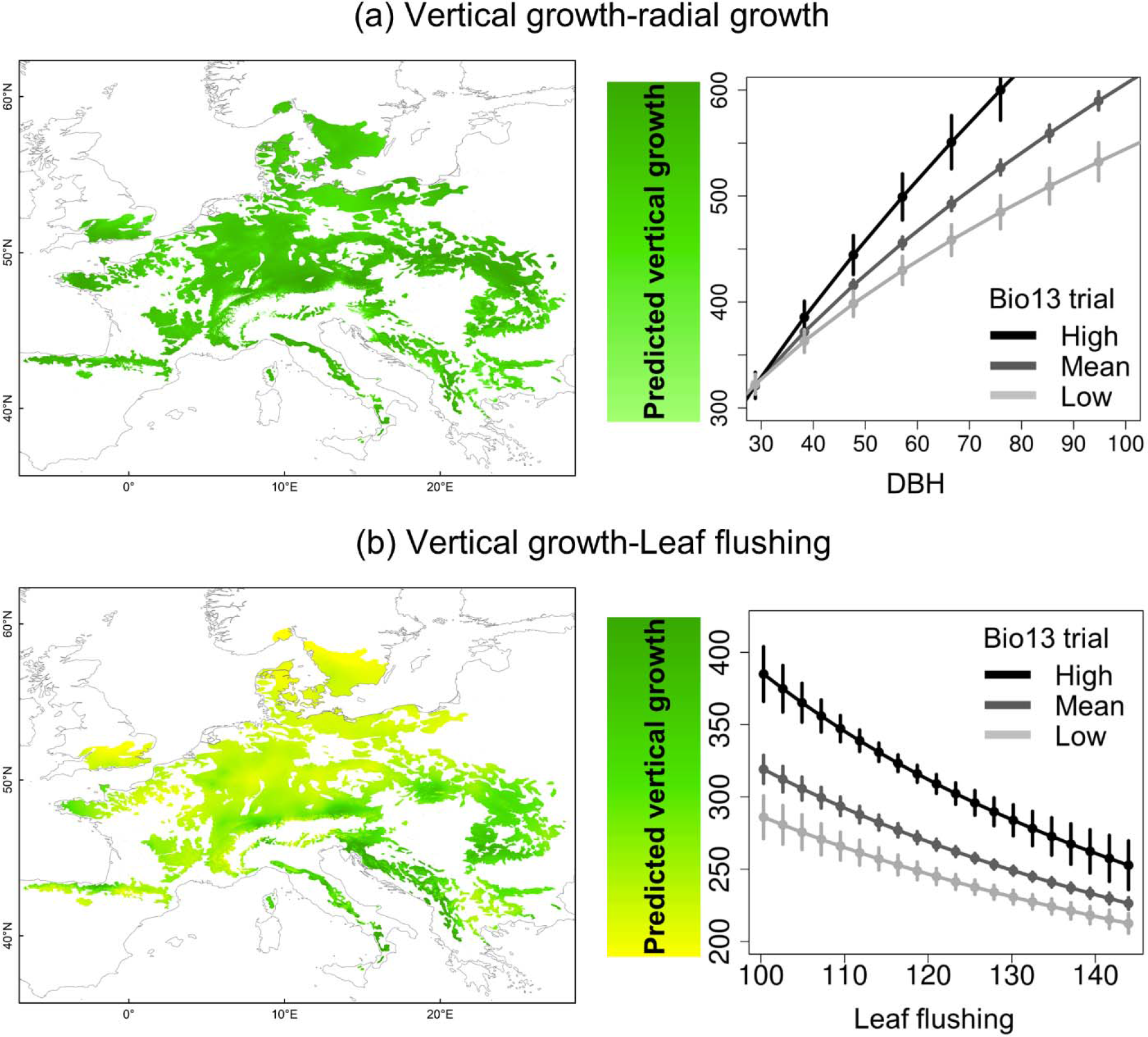
Spatial projections of vertical growth (cm) for (a) Vertical-radial growth model and (b) vertical growth-leaf flushing models (maps on the left), and the corresponding graphs of co-variation between vertical growth and the covariate: (a) DBH (mm) and (b) leaf flushing (Julian days). Black, dark grey, and light grey lines represent high, medium and low values of the precipitation of the wettest month of the trial (BIO13). The vertical lines represent the confidence intervals. The maps display the trait projection for contemporary climate (inferred from 2000-2014 meteorological data) across the current species range. The color gradient depicts the clinal variation in vertical growth from 200 cm (yellow) to 600 cm (green).

### 3.4. Spatial predictions of traits under climate change considering one- and two-trait models

Trait projections for 2070 showed an overall increase in tree growth, particularly for radial growth (Figure 3a, b), but following similar spatial patterns to those predicted under current conditions (Figure 1a, b). Tree survival was predicted to strongly decrease (with respect to that predicted under current conditions, Figure 1c) in the east and throughout the range periphery, while survival rates remained higher in the central part (Figure 3c). Leaf flushing showed similar patterns to those predicted under current conditions (Figure 1d) but with an overall advance in flushing dates (Figure 3d).

**Figure 3.**
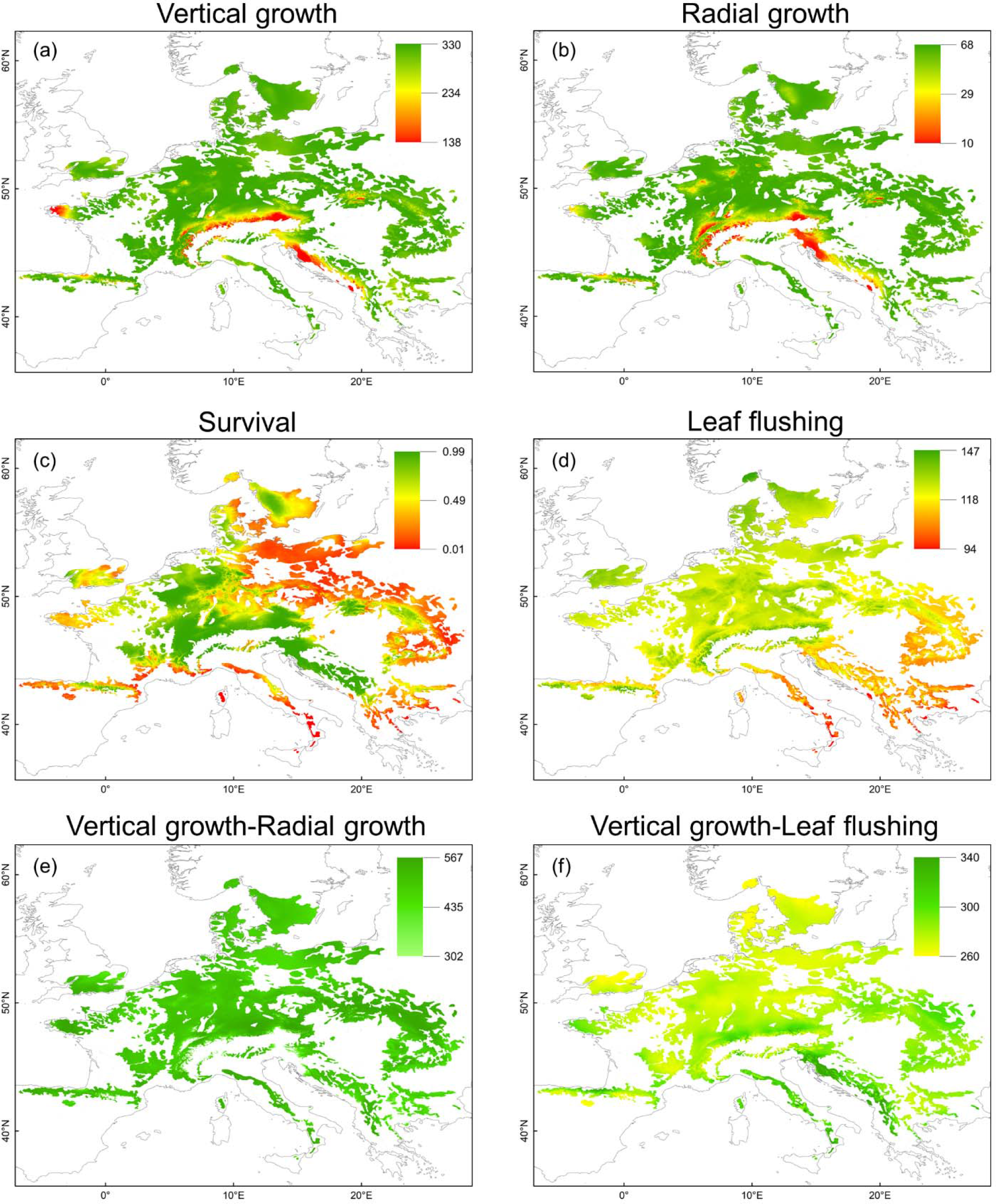
Spatial predictions for 2070 (RCP 8.5) across the species range for one-trait models: (a) vertical growth (cm); (b) radial growth (mm); (c) probability of survival (0=dead; 1=alive); (d) leaf flushing (Julian days); and for two-trait models: (e) vertical growth (cm; co-variate radial growth) and (f) vertical growth (cm; co-variate leaf flushing). The color gradients depict the clinal variation from low (red) to high (green) values.

The prediction of vertical growth, considering radial growth as a covariate, showed an overall increase across the distribution range (Figure 3e) with respect to the model projection of vertical growth under future conditions (Figure 3a). This model showed differences in vertical growth increase compared to the same model applied to current conditions (Figure 2a; Supporting Information Figure S1.3e, Appendix S1). Predictions considering leaf flushing as a co-variate tended to constrain vertical growth throughout the range (Figure 3f) compared with the same model in current conditions (Figure 2b).

### 3.5. Total trait contribution to explain species ranges

All traits and their interactions significantly contributed to explain species occurrence (Table 3). The model explained 31% of the total deviance, with vertical growth accounting for 37%, radial growth for 33%, survival for 19%, and leaf flushing for 1%. The interaction between vertical growth and survival contributed with 3% to the total deviance, that between radial growth and leaf flushing with 2% and the remaining interactions with 1% or less (Table 3).

**Table 3.** Summary statistics for a generalized linear model (binomial family) of beech occurrence (presence/absence) as a function of trait spatial predictions and their interactions. Estimate: coefficient of the regression shown on a logarithmic scale; SE: standard error of fixed variables; *t*: Wald statistical test that measures the point estimate divided by the estimate of its standard error, assuming a Gaussian distribution of observations; *p*: p-value; DE: deviance explained; Vg: vertical growth; Rg: radial growth; S: survival; Lf: leaf flushing.

|  | **Estimate** | **SE** | **$t$** | **$p$** | **DE** |
| --- | --- | --- | --- | --- | --- |
| (Intercept) | -5.84 | 1.15e-02 | -509.03 | 2.00E-16 |  |
| Vg | 5.45 | 1.64e-02 | 332.93 | 2.00E-16 | 0.37 |
| Rg | 0.51 | 7.93e-03 | 64.67 | 2.00E-16 | 0.33 |
| S | 2.11 | 3.75e-03 | 562.83 | 2.00E-16 | 0.19 |
| Lf | 3.12 | 1.48e-02 | 210.94 | 2.00E-16 | 0.01 |
| Vg x S | 0.10 | 4.30e-03 | 21.08 | 2.00E-16 | 0.03 |
| Rg x S | -0.60 | 2.04e-03 | -295.94 | 2.00E-16 | 0.01 |
| S x Lf | -1.40 | 4.02e-03 | -348.1 | 2.00E-16 | 0.01 |
| Vg x Rg | -1.11 | 4.62e-03 | -240.58 | 2.00E-16 | 0.01 |
| Vg x Lf | -7.81 | 2.15e-02 | -363.18 | 2.00E-16 | 0.01 |
| Rg x Lf | 3.43 | 1.09e-02 | 313.89 | 2.00E-16 | 0.02 |
| Model total deviance |  |  |  |  | **0.31** |

## 4. DISCUSSION

### 4.1. Contribution of phenotypic plasticity and local adaptation to range-wide variation in beech growth, survival and leaf flushing

Altogether, our results underpin that range-wide variation in fitness-related traits of beech is driven markedly more by phenotypic plasticity than by local adaptation (Supporting Information Table S1.3, Appendix S1). as happens in other species (Benito Garzón *et al*., *Under review*), and they imply that beech possesses a noteworthy capacity to respond to rapid climate change through acclimation. Although a short-term response through acclimation can be considered as positive for beech to keep pace with climate change, our results point out that the plastic component of tree growth and survival is mostly related to precipitation (Table 2), which follows highly unpredictable patterns (Pflug et al., 2018), making it difficult to evaluate whether acclimation will be enough for beeches to survive (but see our predictions for 2070 under RCP 8.5. showing an increase of mortality in young trees at the margins of the species ranges – Figure 3c). Local adaptation in tree growth (vertical and radial) and survival are driven by adaptation to maximal potential evapotranspiration (Table 2), suggesting that populations are responding to selection factors related to drought (Volaire, 2018). This is in agreement with the general consensus that beech is a drought-sensitive species (Aranda et al., 2015), although there is ongoing debate over the extent of resistance that beech has to drought (Pflug et al., 2018).

The plastic response of leaf flushing to climate was mainly driven by winter temperatures (Table 2). There is a general consensus that these will increase globally in the future (Vautard et al., 2014), and, accordingly, our projection for 2070 anticipates an advance in flushing through most of the range (Figure 1d, 3d and S3d). However, leaf flushing can be constrained by local adaptation to photoperiod (Gauzere et al., 2017; Way & Montgomery, 2015). The fact that phenotypic plasticity and local adaptation in leaf flushing are driven by different environmental parameters implies that these two processes would interact in the long-term. For instance, phenotypic plasticity concerning winter temperatures might enhance local adaptation towards new photoperiodical cues (i.e., shorter spring days), but the evolutionary time scale of local adaptation makes this interaction very unlikely in the short-term.

### 4.2. Trait relationships across the species range

Trait inter-dependence varied to some extent along geographical gradients as the two-trait models had higher predictive power and explained more variance than those based on a single trait (Supporting Information Table S1.3 and Si1.4, Appendix S1). The tight albeit not perfect positive interaction between tree vertical and radial growth (Figure 2a, interaction graph) is unsurprising because of allometric relationships between these two variables, particularly in a common-garden plantation that avoids competition among trees.

The biological basis of the observed co-variation between vertical growth and leaf flushing is less obvious. One possible explanation is that vertical growth is greatly restricted by late flushing in northern beech populations (Kollas, Körner, & Randin, 2014). This would also explain our observation that the one-trait model predicts taller trees to occur in the North, whereas the two-trait model predicts just the opposite. Interestingly, the two-trait model thus implies that strong local adaptation of leaf flushing to photoperiod tends to constrain phenotypic plasticity for vertical growth in northern beech populations (Way & Montgomery, 2015).

### 4.3. Are spatial patterns of growth, survival and leaf flushing delimiting the range of beech?

Beech populations from certain eastern and southern parts of the distribution range seem most sensitive to perturbation, as suggested by the lowest values for all traits considered (Figure 1). Our analysis of species occurrence as a function of spatial trait values also suggests that a combination of these traits contributes differently to the delimitation of the species range (Table 3), in particular: (i) mortality delimits certain parts of the southern and eastern range of beech, reflecting the climatic marginality of the species in these areas, and meaning that these populations are most threatened and making eastwards expansion of beech difficult (although more studies on regeneration are needed to confirm this result); this is the case for many species whose highest mortality is in the driest part of their range (Anderegg et al., 2015; Benito-Garzón et al., 2013; Camarero, Gazol, Sancho-Benages, & Sangüesa-Barreda, 2015); (ii) the smallest girths are predicted in the southern part of the distribution and the eastern part of the range, suggesting that radial growth is mostly restricted by drought (interaction graph and map, Figure 1b), as has already been pointed out (Farahat & Linderholm, 2018); (iii) with very little variation across climatic gradients, vertical growth alone will not delimit beech range. This is not the case for other tree species, for which tree height is clearly delimiting species range (Chakraborty, Schueler, Lexer, & Wang, 2018), highlighting the fact that no single best trait delimits tree species ranges; (iv) projections of trees growing in southern and south-eastern regions that flush early also have higher mortality and lower growth predictions than elsewhere within the species range. However, when tree height and leaf flushing are pooled together in the two-trait model, this leads to an decrease in vertical growth in the North; (v) it seems that in beech, and likely in other species with local adaptation to photoperiod, phenology could restrict the northern expansion of ranges (Duputié et al., 2015; Saltré, Duputié, Gaucherel, & Chuine, 2015), although the link between phenology, survival and fitness is still unclear, and more experiments are needed to better understand the interaction between photoperiod and phenology.

### 4.4. Implications of using trait approaches based on phenotypic variation to forecast beech sensitivity to climate change

Overall, spatial patterns of vertical and radial growth, survival and leaf flushing predicted for the future (Figure 3), are relatively similar to those predicted by the models under current conditions (Figure 1 & 2), which is likely due to the high plasticity of these traits that allows populations to respond to short-term changes in the environment. Our results, based on the study of phenotypic variation, predict species persistence in the future (through high trait values) rather than extinction and migration northwards as predicted by species distribution models based on the occurrence of the species (Kramer et al., 2010; Stojnic et al., 2018).

Nevertheless, the direct comparison of our trait predictions for current and future conditions allows us to detect some differences in their spatial patterns and total trait values (Supporting Information Figure S1.3, Appendix S1), and gives us a better understanding of the temporal dynamics of traits and their relative importance for beech persistence in the future. For instance, our models of leaf flushing predict reduced geographical variability in phenology in the future (from day 94 to 160 -Figure 1d- and from day 94 to 147 – Figure 3d-), as has been reported worldwide (Ma, Huang, Hänninen, & Berninger, 2018). This is mostly explained by larger advances in the phenology of populations at colder sites than those at warmer sites, likely as a consequence of the larger increases in winter temperatures that happen in the North (Kjellström et al., 2018). Survival of young trees is predicted to decrease at the margins of the distribution, but less markedly than is predicted by species distribution models (Kramer et al., 2010; Stojnic et al., 2018).

Including more than one trait related to growth likely reflects a conserved allometric relationship between vertical and radial growth in the future (Figure 3e), but this may be a direct consequence of the lack of competition among trees in our experimental design. Including phenology in two-trait models seems to be detrimental for vertical growth, at least for northern populations where growth is likely constrained by phenology (Figure 3f). However, our trait co-variation approaches are limited to vertical growth as response variables, limiting our understanding of the interplay that other traits can have across species range in the future.

### 4.5. Limitations, perspectives and future research

Although this study relied on the largest network of common gardens for a forest tree in Europe, the resulting inferences suffer from a number of limitations. Our models are based in a limited set of ages (from 2 to 15 years old). However, the expression of phenotypic plasticity changes with age (Mitchell & Bakker, 2014), which can restrict the broad scope of our results to those ages that we considered. This limitation is particularly pronounced for the case of survival (age range 2 to 6 years), for which data only reflect early recruit survival. Other relevant proxies for tree fitness as fecundity and reproduction have not been considered in our approach. In beech, climate warming tends to increase seed production in northern populations (Drobyshev et al., 2010) and to cause a decline in seedling density in southern ones (Barbeta, Peñuelas, Ogaya, & Jump, 2011), which would be expected to continue under climate change.

Our predictions should help to shape future controlled experiments on those populations most sensitive to climate (in the South – East of the range), and others designed to test those trait relationships that are still unclear (phenology – growth – mortality) at the northernmost distribution edge.

## Supporting information

range-wide multi-trait variation

## DATA ACCESSIBILITY

All phenotypic data used in this study are available at https://zenodo.org/record/1240931#.XBuSa81CeUk (Robson *et al.*, 2018). All the maps generated in this study are available from the authors.

## ACKNOWLEDGEMENTS

This study was funded by the “Investments for the Future” program IdEx Bordeaux (ANR-10-IDEX-03-02). HGE was funded by the CONACYT-Mexico and by the Institute of innovation and technology transfer of Nuevo Leon (I2T2), Mexico. TMR was funded by the Academy of Finland (decision 304519). We thank the scientific network INRA-ACCAF.

