## Supplementary material for "Range-wide variation in local adaptation and phenotypic plasticity of fitness-related traits in *Fagus sylvatica* and their implications under climate change": range-wide multi-trait variation

**APPENDIX S1: Supporting information**

**1. Geographical distribution of beech provenance trials**

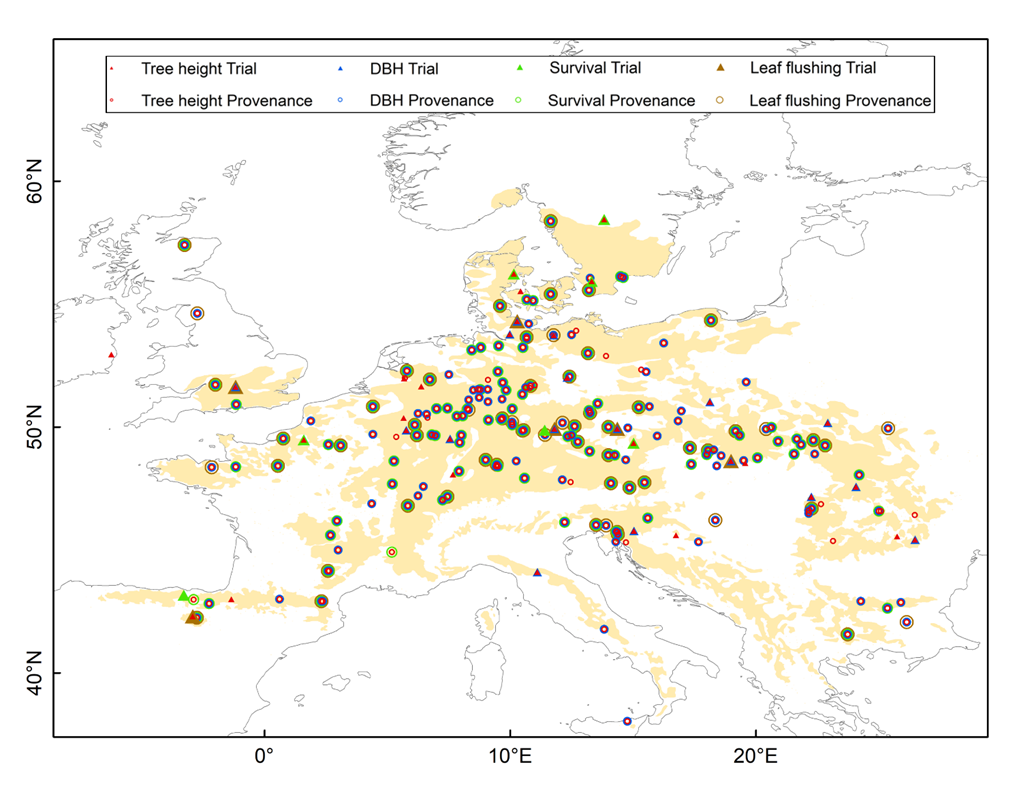

**Supporting Information Figure S1.1.** Distribution range of *Fagus sylvatica* L*.* (shaded in beige) and location of the provenances and trials by trait. Circles indicate the location of the provenances and triangles that of the trials. Different colors have been employed to indicate the different measurements (tree height, DBH, survival and leaf flushing). See Robson et al. (2018) for further details.

**2. Climatic variables**

**Supporting Information Table S1.1.** List of yearly climatic variables provided by EuMedClim. °C: Celsius degree; mm: millimeters; water balance: precipitation minus potential evapotranspiration.

| Climatic variables | Definition | Unit |
| --- | --- | --- |
| BIO1 | Annual mean temperature | °C |
| BIO2 | Mean diurnal temperature range | °C |
| BIO5 | Maximal temperature of the warmest month | °C |
| BIO6 | Minimal temperature of the coldest month | °C |
| BIO12 | Annual precipitation | mm |
| BIO13 | Precipitation of the wettest month | mm |
| BIO14 | Precipitation of the driest month | mm |
| MTdjf | Mean temperature of December, January and February | °C |
| MTmam | Mean temperature of March, April and May | °C |
| MTjaj | Mean temperature of June, July and August | °C |
| MTson | Mean temperature of September, October and November | °C |
| Pdjf | Precipitation of December, January and February | mm |
| Pmam | Precipitation of March, April and May | mm |
| Pjaj | Precipitation of June, July and August | mm |
| Pson | Precipitation of September, October and November | mm |
| Pet Mean | Annual potential evapotranspiration | mm |
| Pet Max | Maximal monthly potential evapotranspiration | mm |
| Pet Min | Minimal monthly potential evapotranspiration | mm |
| Ppet Mean | Annual water balance | mm |
| Ppet Max | Maximal monthly water balance | mm |
| Ppet Min | Minimal monthly water balance | mm |

**3. Principal Components Analysis (PCA) of the climate variables**

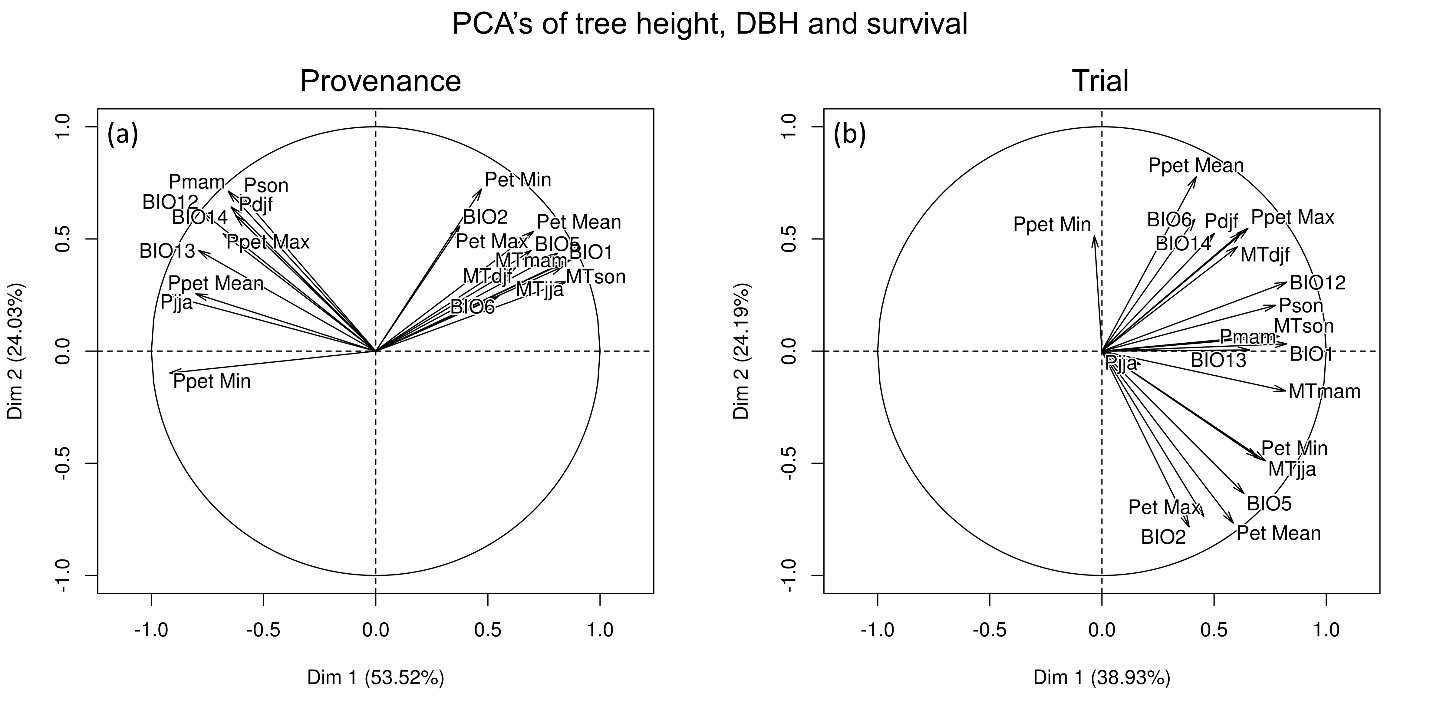

**Supporting Information Figure S1.2.** Results of PCA for selecting climate variables for the models on traits vertical and radial growth, and survival, conducted by provenance (a) and by trial (b). The variance explained by the first two axes is indicated in the figures.

**4. AIC analysis**

We performed a total of 64 one-trait models and selected the best model based on AIC.

**Supporting Information Table S1.2.** AIC values obtained for vertical growth, radial growth, survival and leaf flushing one-trait models. AIC: Akaike information criterion; CP: climate of the provenance; CT: climate of the trial; BIO1: annual mean temperature; BIO5: max temperature of warmest month; BIO6: min temperature of coldest month; BIO12: annual precipitation; BIO13: precipitation of wettest month; BIO14: precipitation of driest month; Pet Max: maximal monthly potential evapotranspiration; Pet Mean: annual potential evapotranspiration; MTdjf: mean temperature of December, January and February; MTmam: mean temperature of March, April and May; MTjja: mean temperature of June, July and August; MTson: mean temperature of September, October and November; MTdjfmam: mean temperature of December, January, February March, April and May.

| **Vertical growth** | | | **Radial growth** | | | **Survival** | | | **Leaf flushing** | | |
| --- | --- | --- | --- | --- | --- | --- | --- | --- | --- | --- | --- |
| **CP** | **CT** | **AIC** | **CP** | **CT** | **AIC** | **CP** | **CT** | **AIC** | **CP** | **CT** | **AIC** |
| Pet Max | BIO13 | 102495.10 | Pet Max | BIO12 | 23099.69 | Pet Max | BIO14 | 39299.61 | MTdjf | MTdjf | -32835.88 |
| BIO13 | BIO13 | 102498.40 | BIO12 | BIO12 | 23099.77 | BIO5 | Pet Max | 39299.75 | MTdjfmam | BIO5 | -32835.2 |
| BIO1 | BIO13 | 102509.20 | Pet Mean | BIO12 | 23100.00 | BIO5 | BIO13 | 39300.20 | MTdjfmam | MTdjf | -32835.01 |
| BIO5 | BIO13 | 102509.70 | BIO5 | BIO12 | 23100.17 | BIO14 | BIO14 | 39300.57 | MTdjf | BIO5 | -32834.71 |
| Pet Mean | BIO13 | 102515.30 | BIO13 | BIO12 | 23105.95 | Pet Mean | Pet Max | 39301.21 | BIO1 | MTdjf | -32833.53 |
| BIO12 | BIO13 | 102538.90 | BIO14 | BIO12 | 23107.76 | Pet Mean | BIO14 | 39303.74 | MTson | MTdjf | -32833.2 |
| BIO6 | BIO13 | 102647.10 | BIO1 | BIO12 | 23109.40 | Pet Max | Pet Max | 39307.04 | BIO1 | BIO5 | -32832.97 |
| BIO14 | BIO13 | 102694.40 | BIO14 | Pet Max | 23112.39 | BIO12 | BIO14 | 39307.83 | MTdjfmam | MTjja | -32832.95 |
| BIO5 | BIO12 | 102827.20 | BIO6 | BIO12 | 23113.15 | BIO5 | BIO12 | 39308.26 | BIO6 | MTdjf | -32832.8 |
| BIO1 | BIO12 | 102836.50 | Pet Max | Pet Max | 23119.66 | BIO13 | BIO13 | 39308.30 | MTdjf | MTjja | -32832.59 |
| Pet Max | BIO12 | 102849.60 | BIO12 | Pet Max | 23119.73 | Pet Mean | BIO1 | 39308.32 | MTson | BIO5 | -32832.53 |
| Pet Mean | BIO12 | 102849.80 | Pet Mean | Pet Max | 23123.73 | BIO5 | BIO14 | 39308.80 | BIO6 | BIO5 | -32831.78 |
| BIO13 | BIO12 | 102856.00 | BIO13 | Pet Max | 23124.58 | Pet Mean | Pet Mean | 39308.84 | BIO1 | MTjja | -32830.75 |
| BIO12 | BIO12 | 102924.80 | BIO5 | Pet Max | 23127.81 | BIO5 | BIO1 | 39308.93 | MTson | MTjja | -32830.39 |
| BIO6 | BIO12 | 103000.30 | BIO6 | Pet Max | 23129.06 | BIO5 | Pet Mean | 39309.13 | MTmam | MTdjf | -32829.82 |
| BIO14 | BIO12 | 103035.00 | BIO1 | Pet Max | 23131.01 | BIO13 | Pet Max | 39310.60 | MTmam | BIO5 | -32829.69 |
| BIO13 | BIO14 | 104366.60 | Pet Mean | BIO13 | 23155.46 | Pet Mean | BIO5 | 39310.84 | BIO6 | MTjja | -32829.67 |
| BIO12 | BIO14 | 104433.70 | BIO1 | BIO13 | 23158.17 | BIO13 | BIO14 | 39311.05 | MTdjfmam | MTson | -32828.93 |
| BIO5 | BIO5 | 104479.60 | BIO5 | BIO13 | 23158.45 | Pet Mean | BIO13 | 39311.74 | MTdjf | MTson | -32828.49 |
| BIO1 | BIO5 | 104486.20 | Pet Max | BIO13 | 23160.20 | Pet Max | BIO13 | 39312.16 | MTmam | MTjja | -32827.31 |
| BIO13 | BIO5 | 104486.40 | BIO6 | BIO13 | 23161.55 | BIO13 | BIO12 | 39312.17 | BIO1 | MTson | -32826.31 |
| Pet Max | BIO5 | 104498.00 | BIO12 | BIO13 | 23170.84 | BIO6 | BIO13 | 39312.88 | BIO5 | BIO5 | -32826.25 |
| Pet Mean | BIO5 | 104502.00 | BIO14 | BIO13 | 23170.94 | Pet Max | BIO1 | 39313.22 | MTson | MTson | -32825.79 |
| BIO12 | BIO5 | 104531.00 | BIO13 | BIO13 | 23172.87 | BIO14 | BIO13 | 39313.52 | BIO6 | MTson | -32825.65 |
| BIO1 | BIO14 | 104548.20 | BIO12 | BIO14 | 23213.00 | BIO5 | BIO6 | 39313.96 | BIO5 | MTdjf | -32825.32 |
| BIO6 | BIO5 | 104551.90 | BIO13 | BIO14 | 23214.59 | BIO12 | Pet Max | 39314.37 | BIO5 | MTjja | -32824.01 |
| Pet Max | BIO14 | 104554.10 | BIO14 | BIO14 | 23221.30 | BIO12 | BIO13 | 39314.57 | MTmam | MTson | -32823.39 |
| Pet Mean | BIO14 | 104561.80 | Pet Max | BIO14 | 23228.03 | BIO13 | BIO1 | 39314.63 | MTdjfmam | BIO1 | -32821.39 |
| BIO5 | BIO14 | 104568.60 | BIO12 | BIO6 | 23228.43 | BIO5 | BIO5 | 39315.15 | MTdjf | BIO1 | -32821.3 |
| BIO14 | BIO5 | 104595.80 | Pet Mean | BIO14 | 23229.18 | Pet Max | Pet Mean | 39315.57 | BIO5 | MTson | -32819.16 |
| BIO14 | BIO14 | 104632.90 | BIO13 | BIO6 | 23230.85 | BIO1 | BIO13 | 39315.98 | BIO1 | BIO1 | -32818.88 |
| BIO6 | BIO14 | 104662.10 | BIO5 | BIO14 | 23231.28 | BIO1 | BIO14 | 39316.04 | MTson | BIO1 | -32818.63 |
| BIO5 | BIO1 | 104948.50 | BIO6 | BIO14 | 23231.86 | BIO6 | BIO14 | 39316.50 | MTjja | BIO5 | -32818.54 |
| Pet Max | BIO1 | 104951.50 | BIO1 | BIO14 | 23235.69 | BIO12 | BIO1 | 39316.56 | BIO6 | BIO1 | -32818.38 |
| Pet Mean | BIO1 | 104953.40 | BIO14 | BIO6 | 23236.45 | Pet Mean | BIO12 | 39316.71 | MTjja | MTdjf | -32817.94 |
| BIO13 | BIO1 | 104958.20 | BIO6 | BIO6 | 23240.51 | BIO6 | BIO12 | 39316.72 | MTjja | MTjja | -32816.41 |
| BIO1 | BIO1 | 104990.00 | Pet Max | BIO6 | 23247.34 | BIO6 | Pet Max | 39316.79 | MTmam | BIO1 | -32815.09 |
| BIO12 | BIO1 | 105034.80 | Pet Mean | BIO6 | 23248.79 | Pet Max | BIO12 | 39316.82 | MTdjfmam | BIO6 | -32813.73 |
| BIO6 | BIO1 | 105103.40 | BIO5 | BIO6 | 23251.22 | BIO13 | Pet Mean | 39317.10 | MTdjf | BIO6 | -32813.64 |
| BIO14 | BIO1 | 105134.00 | BIO1 | BIO6 | 23251.39 | Pet Max | BIO6 | 39317.35 | MTson | BIO6 | -32811.78 |
| BIO13 | Pet Mean | 105607.20 | Pet Max | BIO5 | 23326.60 | BIO1 | Pet Max | 39317.45 | BIO1 | BIO6 | -32811.7 |
| BIO13 | BIO6 | 105655.60 | Pet Mean | BIO5 | 23330.74 | Pet Max | BIO5 | 39317.56 | MTjja | MTson | -32811.33 |
| BIO12 | Pet Mean | 105700.20 | BIO5 | BIO5 | 23333.80 | BIO13 | BIO6 | 39317.60 | BIO6 | BIO6 | -32810.51 |
| BIO12 | BIO6 | 105740.90 | BIO14 | BIO5 | 23336.46 | Pet Mean | BIO6 | 39317.70 | BIO5 | BIO1 | -32810.39 |
| BIO5 | Pet Mean | 105752.70 | BIO12 | BIO5 | 23337.86 | BIO1 | BIO1 | 39317.71 | MTmam | BIO6 | -32807.09 |
| BIO1 | Pet Mean | 105753.10 | BIO13 | BIO5 | 23342.73 | BIO14 | Pet Max | 39317.82 | MTjja | BIO1 | -32803.17 |
| Pet Max | Pet Mean | 105762.50 | BIO1 | BIO5 | 23343.08 | BIO14 | BIO1 | 39317.95 | BIO5 | BIO6 | -32803.03 |
| Pet Mean | Pet Mean | 105769.40 | BIO12 | BIO1 | 23344.66 | BIO14 | BIO12 | 39318.04 | MTdjfmam | MTdjfmam | -32798.69 |
| Pet Max | BIO6 | 105777.20 | Pet Max | BIO1 | 23345.37 | BIO12 | BIO12 | 39318.24 | MTdjf | MTdjfmam | -32798.47 |
| Pet Mean | BIO6 | 105777.90 | BIO6 | BIO5 | 23345.84 | BIO6 | BIO1 | 39318.36 | MTson | MTdjfmam | -32796.38 |
| BIO5 | BIO6 | 105782.20 | BIO5 | BIO1 | 23349.97 | BIO13 | BIO5 | 39318.74 | BIO1 | MTdjfmam | -32796.37 |
| BIO1 | BIO6 | 105790.00 | Pet Mean | BIO1 | 23350.61 | BIO12 | BIO6 | 39320.02 | MTjja | BIO6 | -32795.95 |
| BIO6 | Pet Mean | 105851.70 | BIO14 | BIO1 | 23353.91 | BIO12 | Pet Mean | 39320.05 | BIO6 | MTdjfmam | -32795.67 |
| BIO14 | Pet Mean | 105867.10 | BIO13 | BIO1 | 23354.27 | BIO14 | BIO6 | 39320.34 | MTmam | MTdjfmam | -32792.13 |
| BIO6 | BIO6 | 105898.10 | BIO6 | BIO1 | 23363.77 | BIO6 | BIO6 | 39320.41 | BIO5 | MTdjfmam | -32787 |
| BIO14 | BIO6 | 105901.40 | BIO1 | BIO1 | 23367.18 | BIO12 | BIO5 | 39320.73 | MTdjfmam | MTmam | -32786.71 |
| BIO13 | Pet Max | 106062.80 | BIO14 | Pet mean | 23417.15 | BIO1 | BIO6 | 39321.01 | MTdjf | MTmam | -32785.98 |
| BIO12 | Pet Max | 106132.40 | Pet Max | Pet mean | 23420.69 | BIO1 | BIO12 | 39321.06 | BIO1 | MTmam | -32784.65 |
| BIO1 | Pet Max | 106176.20 | BIO12 | Pet mean | 23423.00 | BIO1 | Pet Mean | 39321.28 | MTson | MTmam | -32784.57 |
| BIO5 | Pet Max | 106179.00 | Pet Mean | Pet mean | 23423.23 | BIO6 | Pet Mean | 39321.81 | BIO6 | MTmam | -32783.62 |
| Pet Max | Pet Max | 106187.20 | BIO5 | Pet mean | 23426.95 | BIO14 | Pet Mean | 39321.88 | MTmam | MTmam | -32780.58 |
| Pet Mean | Pet Max | 106194.00 | BIO13 | Pet mean | 23427.90 | BIO14 | BIO5 | 39322.35 | MTjja | MTdjfmam | -32780.05 |
| BIO14 | Pet Max | 106256.90 | BIO6 | Pet mean | 23431.24 | BIO1 | BIO5 | 39323.48 | BIO5 | MTmam | -32775.52 |
| BIO6 | Pet Max | 106268.70 | BIO1 | Pet mean | 23432.28 | BIO6 | BIO5 | 39323.84 | MTjja | MTmam | -32768.2 |

**5. Summary statistics of one-trait models**

**Supporting Information Table S1.3.** Statistics of random and fixed effects from generalized linear mixed-effect models of vertical growth, radial growth, survival and leaf flushing. Obs: number of trait measurements; Variance: variance explained by the random effects; SD: standard deviation of each level of random effects; Estimate: coefficient of the regression, shown on a logarithmic scale for vertical growth, radial growth and leaf flushing; SE: standard error of each fixed variable; *t*: Wald statistical test that measures the point estimate divided by the estimate of its SE, assuming a Gaussian distribution of observations conditional on fixed and random effects; *z*: Wald statistical test that measures the point estimate divided by the estimate of its SE, assuming a binomial distribution of observations conditional on fixed and random effects. Fixed effects: Coefficients of the fixed effects of the model; CP: climate of the provenance origin; CT: climate of the trial; LatP: latitude of the provenance origin; LatT: latitude of the trial; LongT: longitude of the trial; CP^2^: quadratic effect of the climate of the provenance; CT^2^: quadratic effect of the climate of the trial. Coefficients of the interactions: Age x CP, Age x CT, CP x CT, LatP x CT, LatP x LatT, LatP x LongT, CP x LongT. R^2^M: percentage of the variance explained by the fixed effects (Marginal variance); R^2^C: percentage of the variance explained by the random and fixed effects (Conditional variance); r: Pearson correlation. The climate variable of the provenance (CP) for vertical growth, radial growth and survival is maximal potential evapotranspiration; CP for leaf flushing is mean temperature of December, January and February. The climate variable of the trial (CT) for vertical growth is precipitation of the wettest month, for radial growth is annual precipitation, for survival is precipitation of the driest month and for leaf flushing is mean temperature of December, January and February.

|  | **Vertical growth** | | | **Radial growth** | | | **Survival** | | | **Leaf flushing** | | |
| --- | --- | --- | --- | --- | --- | --- | --- | --- | --- | --- | --- | --- |
| Model | Linear Mixed Effect | | | Linear Mixed Effect | | | Generalized Linear Mixed Effect (Family: binomial) | | | Linear Mixed Effect | | |
|  | Random Effects | | | Random Effects | | | Random Effects | | | Random Effects | | |
|  | Obs | Variance | SD | Obs | Variance | SD | Obs | Variance | SD | Obs | Variance | SD |
| Provenance | 205 | 1.00e-02 | 9.00e-02 | 187 | 9.31e-03 | 9.65e-02 | 114 | 2.98e-01 | 5.46e-01 | 62 | 4.60e-04 | 2.20e-02 |
| Trial | 36 | 9.00e-02 | 3.00e-01 | 19 | 3.81e-01 | 6.17e-01 | 7 | 6.31e-01 | 7.94e-01 | 7 | 3.60e-05 | 6.00e-03 |
| Trial:Block | 107 | 9.00e-02 | 1.00e-01 | 56 | 6.97e-03 | 8.35e-02 | 21 | 1.48e-01 | 3.84e-01 |  |  |  |
| Trial:Block:Tree | 108415 | 8.00e-02 | 2.80e-01 | 31339 | 1.10e-01 | 3.32e-01 | 37433 | 1.16e-02 | 1.08e-01 |  |  |  |
| Residuals |  | 5.00e-02 | 2.20e-01 |  | 1.66e-02 | 1.29e-01 |  | 1.54e-01 | 3.92e-01 |  | 8.56e-04 | 2.92e-02 |
|  | Fixed Effects | | | Fixed Effects | | | Fixed Effects | | | Fixed Effects | | |
|  | Estimate | SE | *t* | Estimate | SE | *t* | Estimate | SE | *z* | Estimate | SE | *t* |
| Intercept | 4.84e+00 | 5.22e-02 | 92.7 | 2.82e+00 | 1.56e-01 | 18.1 | 1.08e+00 | 3.38e-01 | 3.2 | 4.76e+00 | 5.16e-03 | 921.9 |
| Age | 6.45e-01 | 1.14e-03 | 563.6 | 7.17e-01 | 8.74e-03 | 82 | -1.72e+00 | 9.29e-02 | -18.5 |  |  |  |
| CP | 2.58e-02 | 6.93e-03 | 3.7 | 2.94e-02 | 8.81e-03 | 3.3 | 2.83e-02 | 5.30e-02 | 0.1 | 1.07e-02 | 2.63e-03 | 4.1 |
| CT | 9.70e-02 | 4.63e-03 | 20.9 | 2.54e-01 | 7.02e-02 | 3.6 | 1.54e-01 | 2.78e-01 | 0.6 | -1.28e-01 | 9.77e-03 | -13.1 |
| LatP |  |  |  |  |  |  |  |  |  | 5.43e-03 | 2.63e-03 | 2.1 |
| LatT |  |  |  |  |  |  |  |  |  | 4.38e-02 | 4.77e-03 | 9.2 |
| LongT |  |  |  |  |  |  |  |  |  | -1.12e-01 | 9.87e-03 | -11.4 |
| CP2 | -1.27e-02 | 4.84e-03 | -2.6 |  |  |  |  |  |  |  |  |  |
| CT2 | -1.50e-01 | 2.45e-03 | -61.2 | -4.30e-01 | 5.89e-02 | -7.3 |  |  |  |  |  |  |
| Age x CP | -1.07e-02 | 7.86e-04 | -13.6 | -1.09e-02 | 3.58e-03 | 3 |  |  |  |  |  |  |
| Age x CT | -1.92e-02 | 1.50e-03 | -12.8 | 3.33e-01 | 1.44e-02 | 23.1 | 1.59e+00 | 1.21e-01 | 13.1 |  |  |  |
| CP x CT | 9.58e-03 | 1.29e-03 | 7.4 | 7.45e-03 | 3.01e-03 | 2.5 | 8.11e-02 | 2.52e-02 | 3.2 |  |  |  |
| LatP x CT |  |  |  |  |  |  |  |  |  | -1.08e-02 | 1.74e-03 | -6.2 |
| LatP x LatT |  |  |  |  |  |  |  |  |  | 4.15e-03 | 7.95e-04 | 5.2 |
| LatP x LongT |  |  |  |  |  |  |  |  |  | -1.09e-02 | 1.61e-03 | -5.3 |
| CP x LongT |  |  |  |  |  |  |  |  |  | -2.63e-03 | 4.98e-04 | -6.8 |
|  | r | R2 M | R2 C | r | R2 M | R2 C | r | R2 M | R2 C | r | R2 M | R2 C |
|  | 0.69 | 0.57 | 0.91 | 0.53 | 0.51 | 0.98 | 0.59 | 0.18 | 0.40 | 0.73 | 0.49 | 0.68 |

**6. Summary statistics of two-trait models**

**Supporting Information Table S1.4.** Statistics of random and fixed effects from linear mixed-effect models of the vertical growth-radial growth and vertical growth-leaf flushing two-trait models. Obs: number of trait measurements; Variance: variance explained by the random effects; SD: standard deviation of each level of random effects; Estimate: coefficient of the regression shown in logarithmic scale; SE: standard error of each fixed variable; *t*: Wald statistical test that measures the point estimate divided by the estimate of its SE, assuming a Gaussian distribution of observations conditional on fixed and random effects. Coefficients of the fixed effects of the model: Cov: trait covariate; CP: climate of the provenance origin; CT: climate of the trial; CP^2^: quadratic effect of the climate of the provenance. Coefficients of the interactions: Age x CP, CP x CT, Cov x Age and Cov x CT. *R*^2^M: percentage of the variance explained by the fixed effects (Marginal variance); *R*^2^C: percentage of the variance explained by the random and fixed effects (Conditional variance); *r*: Pearson correlation. The trait co-variate (Cov) for growth-radial growth is radial growth and for vertical growth-leaf flushing is leaf flushing. The climate variable of the trial (CT) for the two-trait models is precipitation of the wettest month (BIO13). The climate variable of the provenance (CP) for the two-trait model is maximal potential evapotranspiration.

|  | **Vertical growth-Radial growth** | | | **Vertical growth-Leaf flushing** | | |
| --- | --- | --- | --- | --- | --- | --- |
| Model | Linear Mixed Effect | | | Linear Mixed Effect | | |
|  | Random Effects | | | Random Effects | | |
|  | Obs | Variance | SD | Obs | Variance | SD |
| Provenance | 187 | 1.70e-03 | 4.21e-02 | 150 | 2.33e-02 | 1.53e-01 |
| Trial | 19 | 3.26e-02 | 1.81e-01 | 6 | 1.05e-01 | 3.24e-01 |
| Trial:Block | 56 | 2.20e-03 | 4.60e-02 | 17 | 1.00e-03 | 3.24e-02 |
| Trial:Block:Tree | 31339 | 9.50e-03 | 9.70e-02 | 10634 | 9.82e-02 | 3.13e-01 |
| Residuals |  | 1.50e-02 | 1.23e-01 |  | 2.70e-03 | 5.21e-02 |
|  | Fixed Effects | | | Fixed Effects | | |
|  | Estimate | SE | *t* | Estimate | SE | *t* |
| Intercept | 4.38E+00 | 4.51e-02 | 97.18 | 4.94e+00 | 4.23e-01 | 11.68 |
| Cov | 3.50E-01 | 5.02e-03 | 69.72 | 6.24e-02 | 8.40e-02 | 0.74 |
| Age | -1.97E-01 | 1.26e-02 | -15.66 | 7.40e+00 | 5.28e-01 | 14.01 |
| CP | 5.04E-03 | 3.47e-03 | 1.45 | 2.59e-02 | 1.38e-02 | 1.87 |
| CT | -1.33E-01 | 3.47e-02 | -3.84 | 1.91e+00 | 3.89e-01 | 4.92 |
| CP2 | -5.26E-03 | 2.43e-03 | -2.17 |  |  |  |
| Age x CP |  |  |  | -1.96e-02 | 5.33e-03 | -3.68 |
| CP x CT | -3.47E-02 | 9.66e-03 | -3.59 | 1.78e-02 | 5.72e-03 | 3.11 |
| Cov x Age | 1.05E-01 | 3.44e-03 | 30.57 | -1.43e+00 | 1.09e-01 | -13.08 |
| Cov x CT | 8.02E-02 | 3.82e-03 | 21 | -3.84e-01 | 7.84e-02 | -4.89 |
|  | *r* | *R*2 M | *R*2 C | *r* | *R*2 M | *R*2 C |
|  | 0.76 | 0.62 | 0.95 | 0.77 | 0.47 | 0.99 |

**7. Differences in spatial predictions between future and current climate for one- and two-trait models**

Vertical growth prediction for 12 year-old trees showed small changes in the core of the species range, and moderate decrease in growth in some areas of southern, eastern, western and northern Europe. Increases in vertical growth were mainly expected in the eastern region of the distribution (Supporting Information Figure S1.3a, Appendix S1). Radial growth of 12 year-old trees was predicted to increase in the eastern regions and to decrease across the rest of the range (Supporting Information Figure S1.3b, Appendix S1). Survival of 6 year-old trees was expected to strongly decrease in the western and southern parts of the distribution. Increases in survival were mainly expected in central and some eastern regions of the species range (Supporting Information Figure S1.3c, Appendix S1). The model predicted later leaf flushing in the future than at present for almost all central and western parts of the species distribution. Earlier leaf flushing in the future than today was particularly expected in Sweden (Supporting Information Figure S1.3d, Appendix S1). Differences in vertical growth predictions between future and present climatic conditions (Supporting Information Figure S1.3e and S1.3f). Vertical growth-radial growth models showed an overall increase in vertical growth in some regions of the eastern and southern range; the largest decrease was expected in the southeastern region (Supporting Information Figure S1.3e, Appendix S1). Differences in vertical-growth predictions between the future and present conditions for the vertical growth-leaf flushing model anticipated a decrease in the southeastern and the southern range. A small increase in the northeast was predicted by this model (Supporting Information Figure S1.3f, Appendix S1).

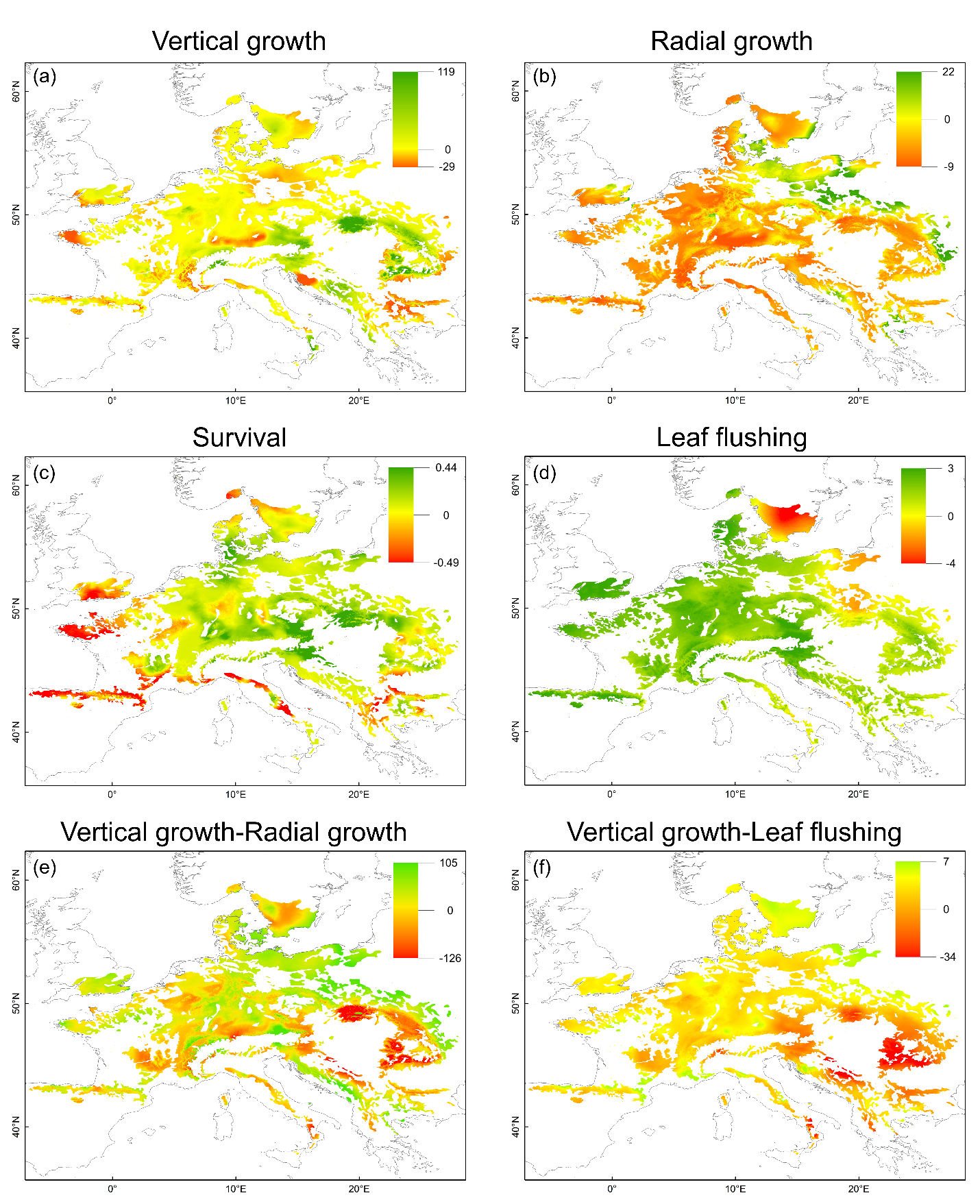

**Supporting Information Figure S1.3.** Differences in predictions between future (2070) and contemporary (2000-2014) climate for one-trait models in beech range: (a) vertical growth of 12 year-old trees (in cm); (b) radial growth of 12 year-old trees (in mm); (c) probability of survival of 6 year-old trees; (d) leaf flushing of 12 year-old trees (difference in Julian days); and for two-trait models: (e) vertical growth (in cm; co-variate radial growth) and (f) vertical growth (in cm; co-variate leaf flushing). The color gradient depicts the clinal variation from low (red) to high (green) values.
